# Late cortical tracking of ignored speech facilitates neural selectivity in acoustically challenging conditions

**DOI:** 10.1101/238642

**Authors:** Lorenz Fiedler, Malte Wöstmann, Sophie K. Herbst, Jonas Obleser

## Abstract

Listening requires selective neural processing of the incoming sound mixture, which in humans is borne out by a surprisingly clean representation of attended-only speech in auditory cortex. How this neural selectivity is achieved even at negative signal-to-noise ratios (SNR) remains unclear. We show that, under such conditions, a late cortical representation (i.e., neural tracking) of the ignored acoustic signal is key to successful separation of attended and distracting talkers (i.e., neural selectivity). We recorded and modelled the electroencephalographic response of 18 participants who attended to one of two simultaneously presented stories, while the SNR between the two talkers varied dynamically. The neural tracking showed an increasing early-to-late attention-biased selectivity. Importantly, acoustically dominant ignored talkers were tracked neurally by late involvement of fronto-parietal regions, which contributed to enhanced neural selectivity. This neural selectivity by way of representing the ignored talker poses a mechanistic neural account of attention under real-life acoustic conditions.

## Introduction

Human listeners comprehend speech surprisingly well in the presence of distracting sound sources (Cherry, 1953). The ubiquitous question is how competing acoustic events capture bottom-up attention (e.g., by being dominant, that is, louder than the background), and how in turn top-down selective attention can overcome this dominance (e.g., listening to a certain talker against varying levels of competing talkers or noise; Kaya and Elhilali, 2017).

Auditory selective neural processing has been mainly attributed to auditory cortex regions. It is by now well-established that the auditory cortical system selectively represents the (spectro-)temporal envelope of attended, but not ignored speech (i.e., neural phase-locking; Magneto-encephalography: Ding and Simon, 2012; Electroencephalography: Kerlin at al., 2010; Power et al., 2012; Horton et al., 2013; O’Sullivan et al., 2014). Accordingly, auditory cortical responses allow for a reconstruction of the spectrogram of speech and to detect the attended talker (e.g., Mesgarani and Chang, 2012; Zion Golumbic et al., 2013). In sum, selective neural processing in auditory cortices establishes an isolated and distraction-invariant spectro-temporal representation of the attended talker.

However, as has been shown, degradations of the acoustic signals attenuate the neural phase-locking to speech. Experimental degradations have included artificial transformations of temporal fine structure (Ding et al., 2014; Kong et al., 2015), or rhythmicity (Kayser et al., 2015), reverberation (Fuglsang et al., 2017) or decreased signal-to-noise ratio (SNR; Kong et al., 2014; Ding and Simon, 2013; Giordano et al., 2017). Not least, neural selection of speech appears weakened in people with hearing loss (Petersen et al., 2016). In sum, those studies suggest that the strength of neural phase-locking indicates behavioral performance such as speech comprehension.

Additionally, higher order non-auditory neural mechanisms facilitate speech comprehension as well. The supra-modal, fronto-parietal attention network is a candidate to be involved in top-down selective neural processing during demanding listening tasks (Woolgar et al._(_ 2016). Beyond the phase-locking in lower frequency bands (i.e., ~1 −8 Hz; Wang et al 2018, Pomper and Chait 2017), top-down selective neural processing has also been associated with changes in the power of induced alpha-oscillations (i.e., ~8 - 12 Hz; Obleser and Weisz 2012; Kayser et al. 2015, Wöstmann et al. 2016). Specifically, increased parietal alpha-power is related to enhanced suppression of the distracting input (Wöstmann et al., 2017). This reflects that, besides the neural spectro-temporal enhancement of the attended talker, a crucial role in top-down neural selective processing was attributed to the suppression of the ignored talker.

Neural signatures of suppression can be two-fold. First, suppression can attenuate the neural response to an ignored talker compared to an attended talker, like it was found in neural phase-locking from latencies of around 100 ms (Ding and Simon, 2012; Wang et al., 2018). Second, active suppression can add or increase components in the neural response to the ignored talker, given that the response is dissociable from the response to the attended talker (e.g.; a louder ignored talker evoking a stronger neural response anti-polar to the response to a louder attended talker). Here we asked, how the components of the phase-locked neural response are affected by selective attention under varying signal-to-noise ratio (SNR).

The phase-locked neural response to broad-band continuous speech can be obtained from EEG by estimating the (delayed) covariance of the temporal speech envelope and the EEG, which results in a linear model of the cortical response; a temporal response function (TRF; Lalor et al., 2009; Crosse et al., 2016). Analogous to the event-related potential (ERP), the components of the TRF can be interpreted as reflecting a sequence of neural processing stages where later components reflect higher order processes within the hierarchy of the auditory system (Davis and Johnsrude, 2003; Picton et al., 2013; Di Liberto et al., 2015).

Here, we use a listening scenario in which two concurrent talkers undergo continuous SNR variation. Our results demonstrate differential effects of bottom-up acoustics vs. top-down selective neural processing on earlier vs. later neural response components, respectively. Source localization reveals that not only auditory cortex regions are involved in the selective neural processing of concurrent speech, but that a fronto-parietal attention network contributes to selective neural processing through late suppression of the ignored talker.

**Figure 1:**
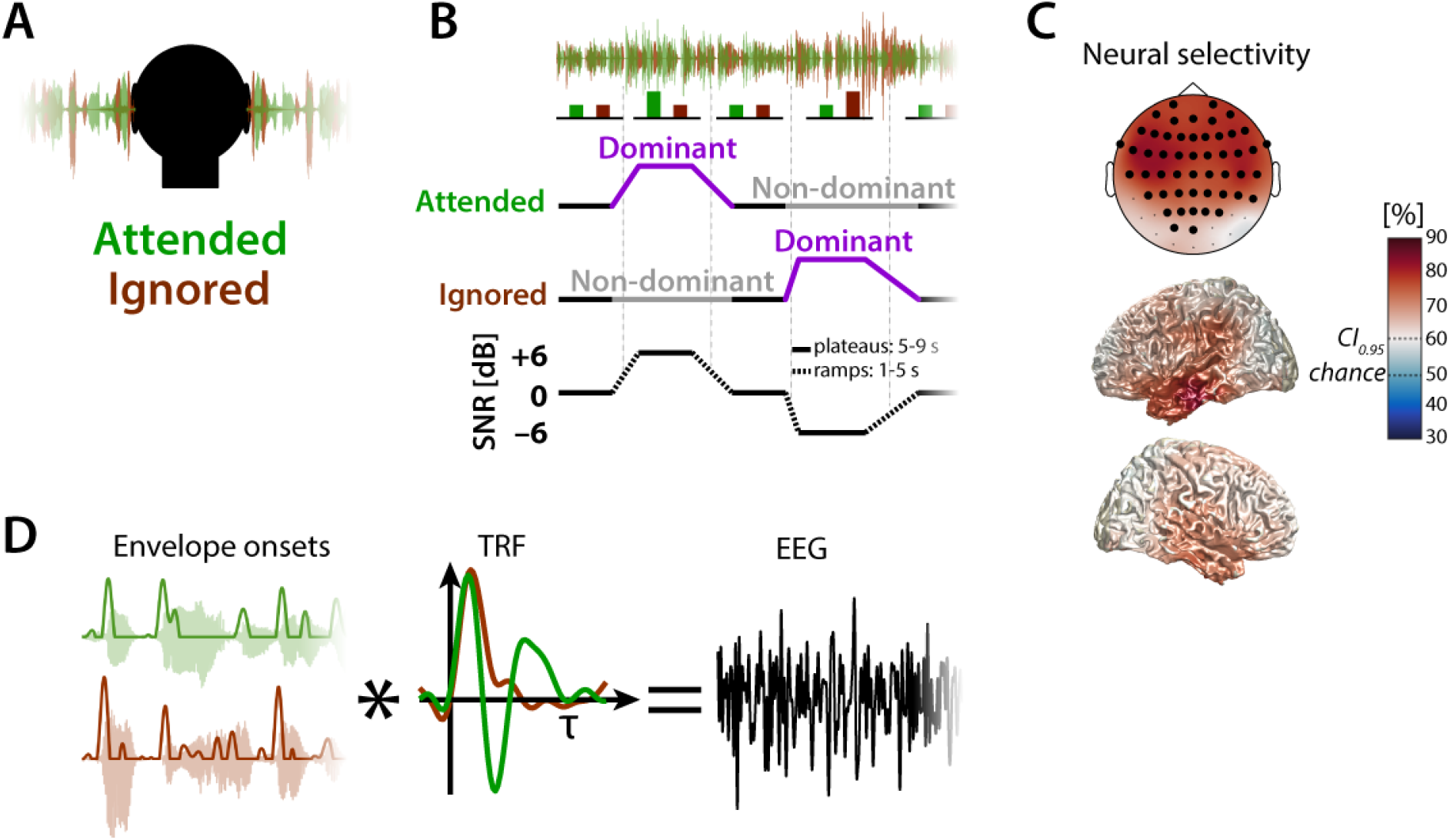
Experimental design, forward model, and neural selectivity. **A)** Two mixed talkers (female & male) were presented on both ears without spatial segregation (diotic). **B)**The signal-to-noise ratio (SNR) between attended (signal) and ignored (noise) talker was varied between −6, 0 and +6 dB by either raising the level of the attended talker or the ignored talker. Length of ramps and plateaus were drawn from uniform distributions. **C)**Neural selectivity here expressed as classification accuracy in detection of the attended and ignored talker averaged across subjects. Shown here is accuracy as obtained by prediction of EEG signals (Fiedler et al., 2017) at single EEG channels and single voxels in source space, respectively. Highlighted channels of topographic maps indicate that the lower bound of the confidence interval (bootstrapped mean on the group level) was greater than the 95%-confidence bound of a binomial distribution (Cl_0.95_ = 60%). **D)**Temporal response functions (TRF) to the attended and ignored talker were extracted by a forward (encoding) regression model based on the assumption that the measured EEG signal is the superposition (convolution) of the envelope onsets (of the attended and ignored talkers) and the TRFs, respectively. TRFs reflect the neural response evoked by a single envelope onset.

## Results

We asked participants to listen to one of two simultaneously presented audiobooks under varying signal-to-noise ratio (Fig. 1A&B; −6 to +6 dB SNR). After each of twelve five-minute blocks, subjects were asked to rate the difficulty of listening to the to-be-attended talker on a color bar ranging from red (difficult = 1) to green (easy = 10). The average difficulty ratings strongly varied between subjects (mean: 5.2, SD: 2.2, range: 2.3-8.9). No difference in difficulty ratings for listening to the female versus the male talker was found (one-sample t-test, t_17_ = 1.17, p = 0.26).

To test their successful attending, participants were asked to answer four multiple-choice questions on the content of the to-be-attended audiobook after each five-minute block. The percentage of correctly answered questions was far above chance (25%) for all participants (mean: 81 %, SEM: 2%, range: 60–96%). All participants were thus able to follow the to-be-attended talker.

### Neural selectivity

To obtain a general estimate of which EEG channels and which voxels reveal signatures of *neural selectivity*, we identified the attended (and the ignored) talker by forward prediction of EEG signals based on one-minute parts of the EEG and envelope onsets (see methods). Overall *neural selectivity* was highest (up to 80%) at fronto-central electrodes and respective temporal cortex regions in source-space (Fig. 1C).

**Figure 2:**
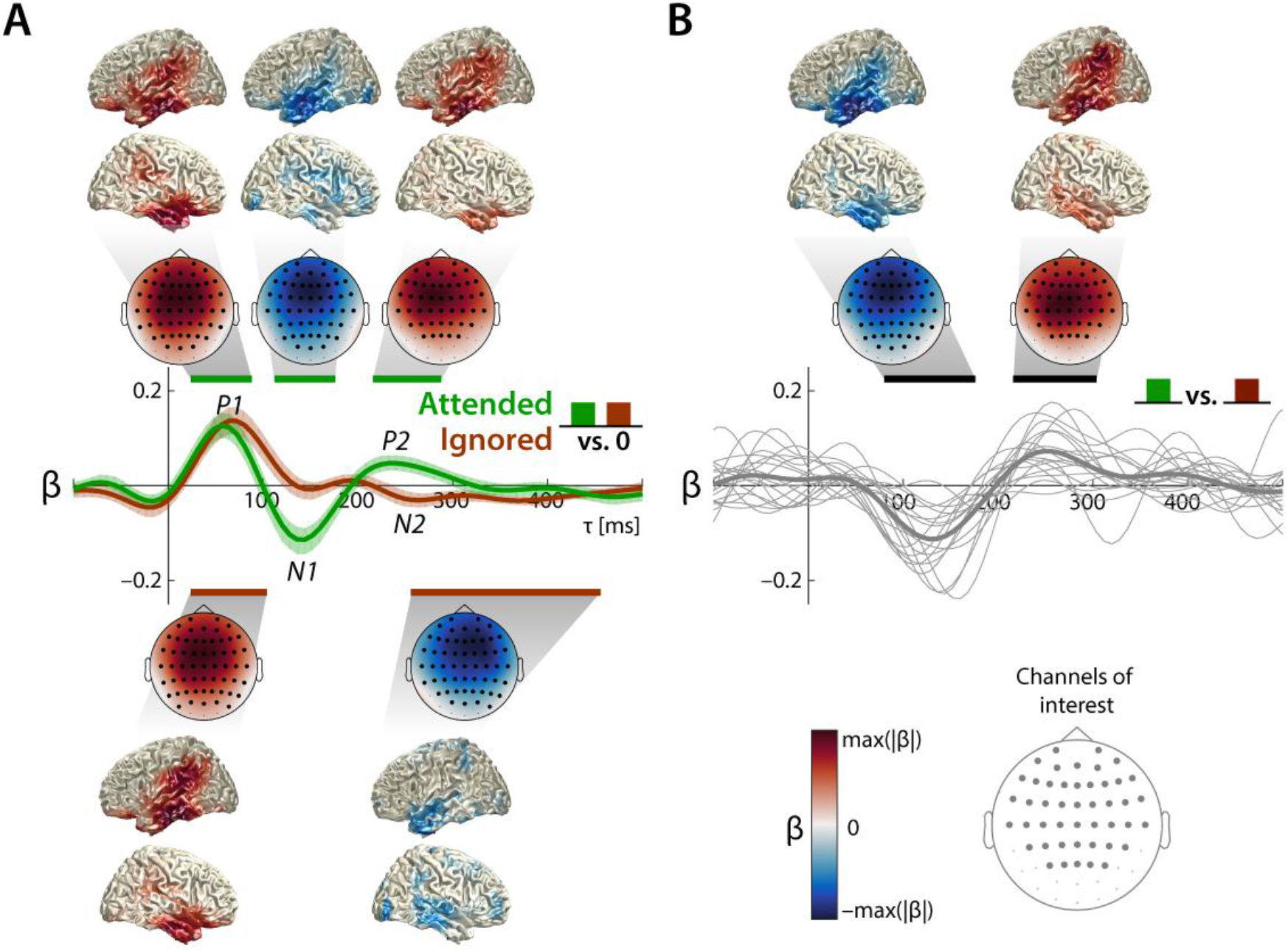
Temporal response functions (TRF) to continuous speech of concurrent talkers under balanced SNR (0 dB). TRF β-weights depict average across subjects and average across channels of interest. Confidence bands (95%) were obtained by bootstrapping the mean across subjects. Horizontal lines indicate time ranges of significant difference from zero obtained from a cluster-based permutation test at the group level. Topographic maps showβ-weights of clusters averaged across the cluster time range. Highlighted channels are part of the significant clusters. Source localizations show the 20% most strongly contributing voxels. **A)**Response to the attended talker (green, upper topographic maps) clearly show a cascade of three components (P1_TRF_-N1_TRF_-P2_TRF_).Response to the ignored talker (red, lower topographic maps) only show a P1_TRF_, whereas the N1_TRF_ and P2_TRF_are suppressed. **B)**Significant differences between neural responses to the attended and ignored talker are present in the N1_TRF_-and P2_TRF_-timerange. Thin grey lines show single subject TRFs averaged across channels of interest.

### Attention modulates neural responses to concurrent speech

Next, we assessed in greater detail the unfolding of attentional selection of to-be-attended speech in time. To this end, we estimated the TRFs from the balanced SNR trials of 0 dB (i.e. independent of the SNR manipulation) and assessed the most prominent response components and their modulation by attention. We inspected both the TRFs to the attended and ignored talker individually (Fig. 2A), as well as the difference between the TRFs to the attended and ignored talker (Fig. 2B) to examine signatures of *neural selectivity*.

First, an early positive component (termed P1_TRF_) appeared in the TRFs to the attended (Fig. 2A, 24-88 ms, p = 2×10^4^) and ignored (Fig. 2A, 24-112 ms, p = 2×10^4^) talkers, but without any attention-related difference (Fig. 2B). Latency, polarity, and topography of this component compared well to a P1 as found in auditory evoked potentials (AEPs).

Second, a later negative deflection (termed N1_TRF_) was only present in the TRF to the attended talker (Fig. 2A; 112-176 ms, p = 5×10^−4^). This component was significantly increased in magnitude (i.e., more negative) for the attended versus the ignored talker (Fig. 2B, 80-176 ms, p = 5×10^−4^; see also Fig. S3). Noteworthy, the significant attentional modulation of this component (attended-ignored) started already at a time lag of 80 ms, when both the TRF to the attended and to the ignored talkers were still in positive deflection (see Fig. 2A).

Third, a positive deflection between 200 and 300 ms (termed P2_TRF_; Fig. 2A, 216-304 ms, p =5×10^−4^), was again only present in the TRF to the attended talker. This component mainly drove the significant difference between the responses to the attended and ignored talker (Fig. 2B, p =2×10^−4^).

Interestingly, in the same time interval, a negative deflection was found in the TRF to the ignored talker (termed N2_TRF_; Fig.2B, 248-424 ms, p = 2×10^−4^). While at earlier stages, TRFs to the attended and the ignored talker showed the same polarity (P1_TRF_), at the stage of the P2_TRF_ we see an anti-polar relationship. Effectively, this also enhanced the late, attended-ignored difference in the P2_TRF_ time range (Fig. 2B).

In sum, three prominent components (P1_TRF_, N1_TRF_, P2_TRF_; Fig. 2A) were identifiable with notable consistency across individual subjects. The latter two components were absent in the TRF to the ignored talker and thus indicated *neural selectivity*. All three components (P1_TRF_, N1_TRF_, P2_TRF_) mainly localized to superior and inferior temporal regions (Fig. 2A). Note that the source localizations of the two latter components (N1_TRF_, P2_TRF_) compared well to the sources of enhanced neural selectivity between attended and un-attended talkers (Fig. 1C).

### Late representation of ignored talker enhances towards more detrimental SNRs

Next, we analyzed the impact of a varying SNR on the Temporal response functions (TRFs). To this end, we first contrasted the TRFs of the two extreme conditions (SNRs −6 vs. +6 dB; Fig. 3A&B). Second, we contrasted TRFs across SNRs matched for the acoustic properties of being either the louder or the quieter talker (Fig. 3C&D), such that the occurring differences between the TRFs to the attended and the ignored talker can solely be related to top-down attending versus ignoring. For simplicity, we will use the terms *dominant* (attended talker under +6 dB SNR, ignored talker under −6 dB SNR) and *non-dominant* (attended talker under −6 dB SNR, ignored talker under +6 dB SNR). We observed an SNR-dependent latency shift which hindered time-lag-wise attended-ignored contrasts within SNRs (Fig. 3A&B, see appendix for more details).

Importantly, two later additional components appeared whenever the ignored talker was dominant (Fig. 3B): the first (160-178 ms, p = 0.04) localized to temporal regions, while the second extended markedly into parietal regions (232-280 ms, p = 0.001). The enhanced involvement of parietal regions differentiated this detrimental-SNR, ignored-speech component from all others. Visual inspection of the TRFs to dominant talkers (Fig. 3C) highlights the additional late N2 component in the TRF to the ignored talker, which appears to be anti-polar to the P2_TRF_ to the attended talker.

In contrast, TRFs to *non-dominant* talkers (Fig. 3D) suggest that the observed attention-related differences are decreased (cf., Fig. 3C) due to smaller deflections of the N1_TRF_ and P2_TRF_ to the *non-dominant* attended talker and the lack of the anti-polar N2_TRF_ to the *non-dominant* ignored talker. We summed the magnitude of the attended-ignored difference across all time lags, which revealed a smaller attended-ignored difference for *non-dominant versus dominant* talkers (t_17_ = 3.80, p = 0.0014). Thus, the neural response to a *dominant* ignored talker does not resemble the neural response to a dominant attended talker by capturing bottom-up attention. Instead, dominant ignored speech retains a distinct “ignored” neural signature, most likely to due to top-down neural signaling of its to-be-ignored status.

In sum, our findings indicate that, when a talker is *dominant*, neural signatures of selective processing are enhanced (compared to *nondominant*). Importantly, this enhancement is not only affecting the representation of the attended talker, but an important contribution to this enhanced top-down processing can be attributed to an additional late component (N2_TRF_) in the neural response to the ignored talker. To further disentangle the contribution of the selective processing of the attended and ignored talker, we established the time lag and talker resolved measures *neural tracking* and *neural selectivity*, which will be discussed in the following section.

**Figure 3:**
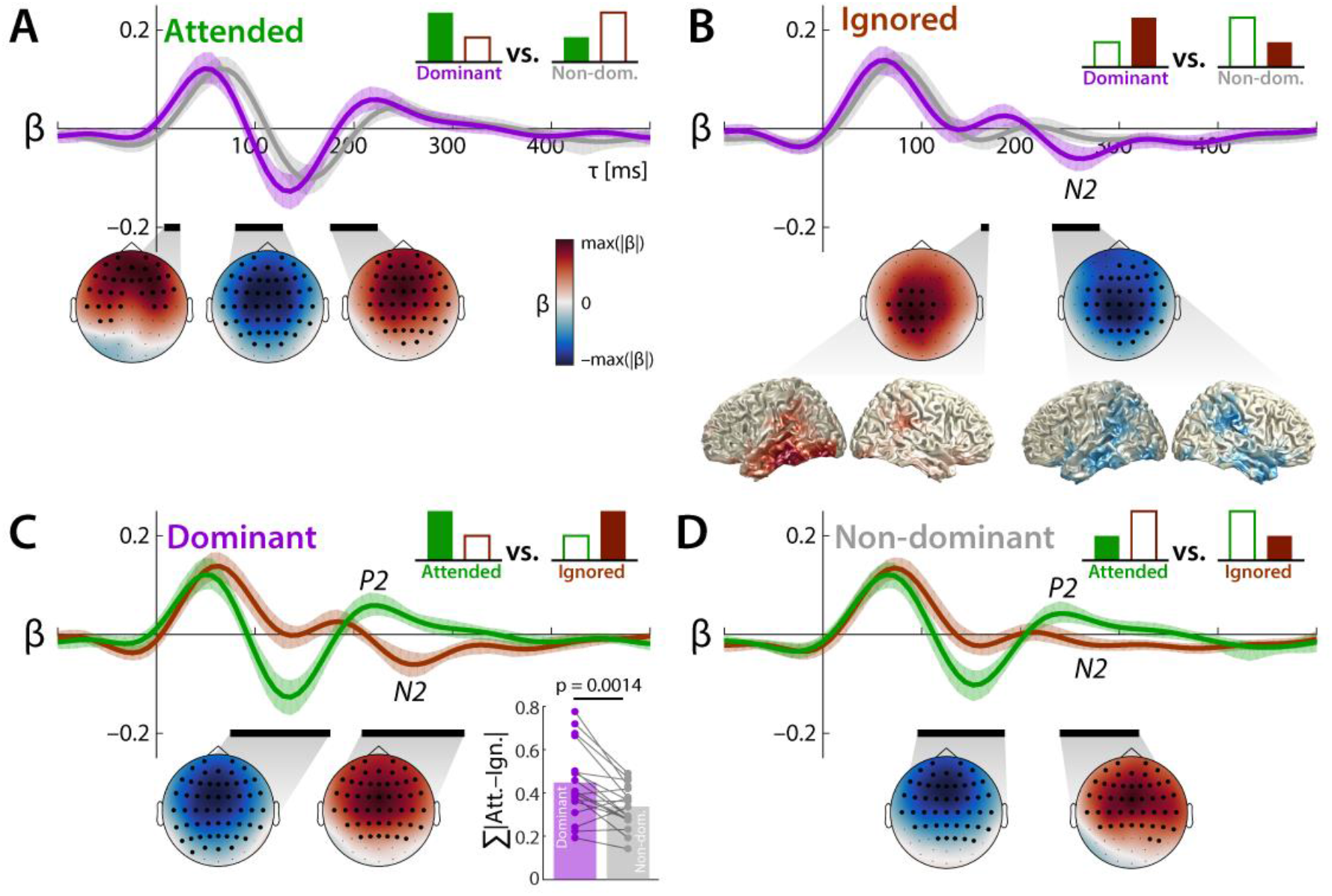
Temporal response functions (TRF) to continuous speech of concurrent talkers contrasted as dominant vs. non-dominant talkers and attended vs. ignored talkers, respectively. TRF β-weights depict average across (N = 18) subjects and average across channels of interest. Confidence bands (95%) were obtained by bootstrapping the mean across subjects. Schematic bar graphs indicate the investigated contrast. Black horizontal lines indicate time ranges of significant difference obtained from a cluster-based permutation test at the group level. Topographic maps showβ-weight differences of clusters averaged across the cluster time range. Highlighted channels are part of the significant clusters. Source localizations show the 20% most strongly contributing voxels with full opacity. **A)** Responses to the non-dominant attended talker are delayed compared to the dominant attended talker. **B)** A late component appeared in the response to the dominant ignored talker, which involved parietal regions. **C)** Late negative response (N2_TRF_) to the dominant ignored talker appears anti-polar to the response to the dominant attended talker. Inset: Magnitude of the attended-ignored TRF difference summed across all time lags for *dominant* and *non-dominant* talkers. **D)** Non-dominant talkers show significant but decreased attention-related differences.

### Neural selectivity increases by way of a late cortical representation of ignored speech

We established two measures to quantify the encoding and the selective neural processing of the talkers during the unfolding of the neural response reflected in the TRFs. First, *neural tracking* is a measure of how strongly a single talker is represented (i.e., encoded) in the EEG. Second, *neural selectivity* quantifies how accurately an attended talker can be identified as attended and an ignored talker as ignored, respectively.

Parallel inspection of *neural tracking* and *neural selectivity*allowed us to disentangle the effects of bottom-up and top-down attention on the TRFs. For example, the increased sound pressure level of a talker may increase its saliency and thus bottom-up pull attention towards it. This would result in enhanced *neural tracking* of the ignored talker and the neural response would become less distinct from the respective response to a *dominant*, but intentionally attended talker. However, if there exists a counter-acting, top-down process that enhances and maintains a neural-response differentiation between the attended and the ignored talker, *neural selectivity* would increase at the same time.

To get a total estimate of *neural tracking* of the two talkers, we first used all time lags of the TRFs (i.e., −100-500 ms). Fig 4A shows the *neural tracking* of the attended, the ignored as well as the overall tracking of the two talkers (attended & ignored). The overall tracking was found to be well above zero for all participants as well as the tracking of the two talkers separately (Fig. 4A, bottom).

In a next step, we estimated the time-lag- and channel-dependent unfolding of *neural tracking*. As expected, we found enhanced *neural tracking* of the attended talker compared to the ignored talker between 144 and 288 ms under the balanced SNR of 0 dB (Fig. S1 A), driven by fronto-central channels. This is congruent with the time range and topography of the N1 TRF and P2_TRF_, which were found to be non-present in the TRF to the ignored talker.

Interestingly, towards more adverse SNRs (dominant ignored talker), the late enhanced *neural tracking* of the attended talker compared to the ignored talker seems to shrink (Fig. 4B). Visual inspection of the time-lag resolved *neural tracking* suggests that this shrinkage is due to an additional late cortical representation of the ignored talker that appears whenever the ignored talker is *dominant*. The contrast of the *neural tracking* of the dominant and the non-dominant ignored talker confirmed such a late cortical representation (Fig. 4C, 240-312 ms, p = 1.5×10^−3^) originating mainly from fronto-parietal as well as temporal regions.

Importantly, the overall *neural selectivity* is not affected by adverse conditions (Fig. 4E, grey bars, −6 vs +6 dB, one-sample t-test, t_17_= 0.24, p = 0.81). However, the relative contribution of the neural selectivity of the attended talker and ignored talker changes across SNRs (−6 vs +6 dB; one-sample t-test; attended: t_17_= −4.6, p = 2.77×10^−4^; ignored: t_17_= 2.18, p = 0.044): Towards more adverse SNRs, the *neural selectivity* of the ignored talker increases, while the *neural selectivity* of the attended talker decreases (Fig. 4E, top). This is also discernible in single subjects (Fig. 4E, bottom), where *neural selectivity* of the attended talker is stronger under an SNR of +6 dB (right, 16 of 18 subjects) and stronger for the ignored talker under an SNR of-6 dB (left, 11 of 18 subjects).

If the increased *neural tracking* of the dominant ignored talker at later stages (Fig. 4C) is solely driven by its increased saliency (i.e., higher dominance evoking a stronger response), we would expect no concomitant increase in neural selectivity (see above). However, we found a late increase in *neural selectivity* for the dominant compared to the *non-dominant* ignored talker (Fig. 4G, 216-264 ms, 2.5×10 ^3^). Neural sources compared well to the increased fronto-parietal *neural tracking* of the dominant ignored talker (see Fig.4C&G).

Further more, *neural tracking* and *neural selectivity* (for *dominant* vs *nondominant* ignored speech) were positively correlated (Fig. 4D, r = 0.78, p = 0.014×10^−2^): If a listener’s neural tracking was relatively strong for the dominant versus non-dominant ignored talker, the neural response allowed more accurate identification of the ignored talker as ignored.

In sum, at later stages, not only increased selective neural processing of the attended talker but also the selective neural processing of the ignored talker facilitates input segregation under adverse listening conditions.

**Figure 4:**
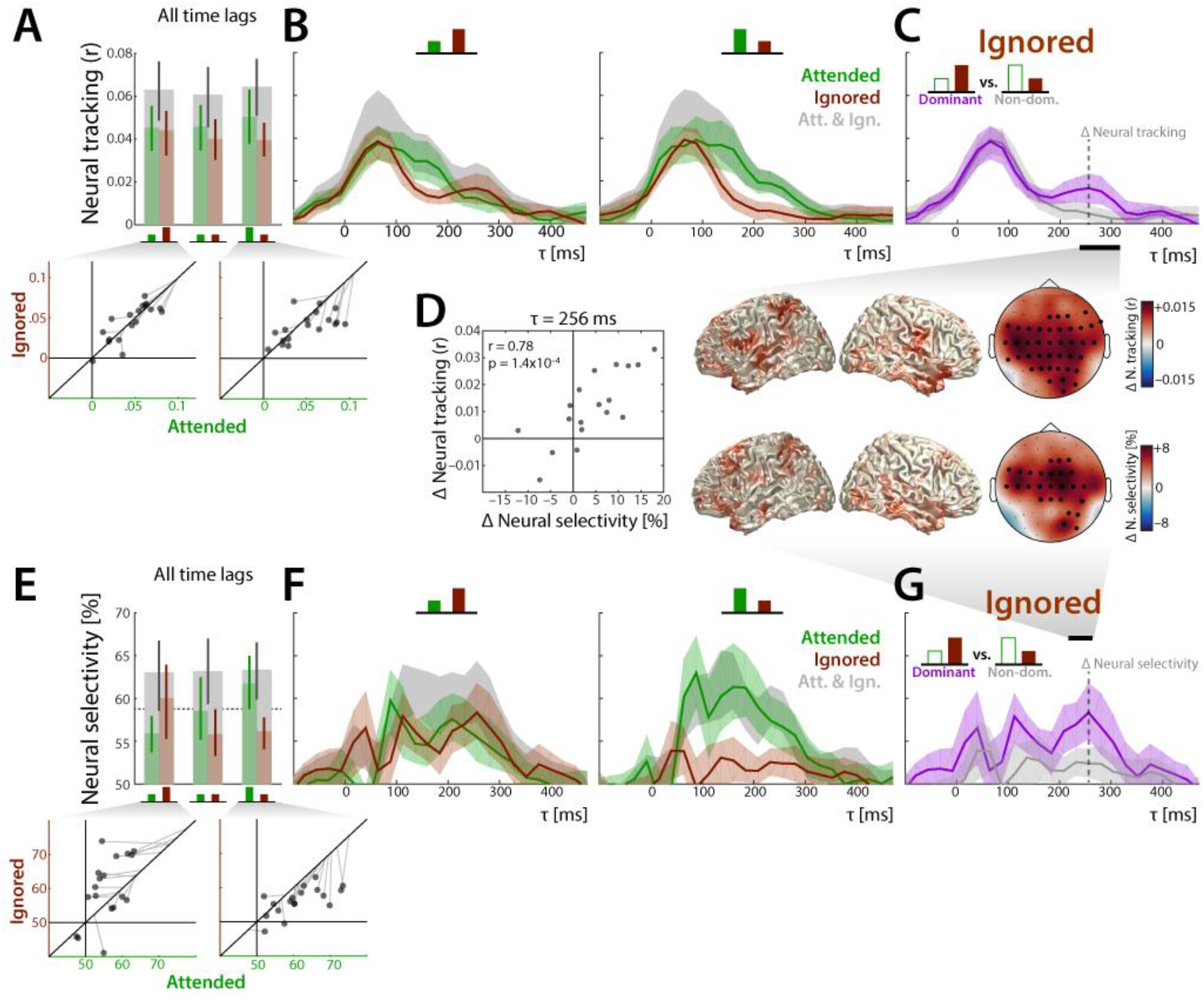
Unfolding of neural tracking and neural selectivity reveals late neural selective processing of the ignored talker. Neural tracking and neural selectivity were estimated based on the extracted TRFs to the attended (green), the ignored (red), as well as both talkers (grey). Confidence bands (95%) were obtained by bootstrapping. Highlighted channels (topographic maps) are part of a significant cluster. Source localizations show the 20% most strongly contributing voxels with full opacity. **A)** Neural tracking across all time lags (−100-500 ms). Scatterplots (bottom) show single-subject data averaged across channels of interest. Grey lines indicate overall neural tracking of both talkers at the45°-line. **B)** Unfolding of neural tracking across time lags under SNR of-6 (left) and +6 dB (right). **C)** Contrast of neural tracking between the dominant and non-dominant ignored talker. **D)** Correlation of change in neural tracking and change of neural selectivity at τ= 256 ms. **E)** Neural selectivity across all time lags (−100-500 ms). Scatterplots (bottom) show single-subject data averaged across channels of interest. Grey lines indicate overall neural tracking of both talkers at the 45°-line. **F)** Unfolding of neural selectivity across time lags under SNR of-6 (left) and +6 dB (right). **G)** Contrast of neural selectivity between the dominant and non-dominant ignored talker.

## Discussion

In the present study, human listeners attended to one of two concurrent talkers under continuously varying signal-to-noise ratio (SNR). We asked to what extent a late cortical representation (i.e., neural tracking) of the ignored acoustic signal is key to the successful separation of to-be-attended and distracting talkers (i.e., neural selectivity) under such demanding listening conditions.

Forward modeling of the EEG response revealed neural responses to the temporal envelopes of individual talkers and their modulation by both, top-down attentional set, and bottom-up SNR. Critically, towards more adverse SNRs, an additional late negative component occurred in the neural response to the ignored talker. Under adverse conditions, this component was found to be accompanied by enhanced selective neural processing *(neuralselectivity)*, emerging primarly from fronto-parietal brain regions.

The present result suggests that irrelevant, to-be-ignored acoustic inputs are not simply absent from the late cortical response but become actively suppressed in regions beyond auditory cortex.

### Early and late neural signatures of selective neural processing

Generally, we replicated previous results that showed that attention-ignored differences in the neural response can mainly be found at time lags > 80 ms, which were mainly attributed to stronger neural tracking caused by enhanced N1 and P2 components in the response to the attended vs. ignored talker (Horton et al. 2013; O’Sullivan et al., 2014; Ding & Simon, 2012). Here we show that a P2-counter-acting response to the ignored talker enhances the attended-ignored difference as well.

While earlier studies showed that selective neural processing in auditory cortices is mainly working out a clean representation of the attended talker (Mesgarani and Chang, 2012; Zion Golumbic, 2013), we show that a late neural representation of a distracting auditory input is accompanied with enhanced selective neural processing in a cocktail-party scenario as well. This additional late neural representation was revealed by going beyond strictly matched sound pressure levels of attended versus ignored speech (cf., Horton et al., 2013; O’Sullivan et al., 2014, Ding & Simon, 2012; Mirkovic et al., 2015; Biesmanns et al., 2016), by presenting speech signals both as target and distractor (cf., Ding & Simon, 2013) and by applying SNR-variation symmetrically around 0 dB (cf., Kong et al., 2014). In sum, our design allowed us to draw conclusions on the neural selective processing of real-world listening scenarios of dynamically varying listening demand.

Our investigation of concurrent speech under varying SNR helps disentangle neural mechanisms of early and late selection (Treisman 1964). Since the ignored talker predominantly masks the attended talker under adverse listening conditions (i.e., negative SNRs, which we have labelled *dominant*), early neural filters tuned to the spectro-temporal properties of the attended talker might not be sufficient (i.e., neural gain, Willmore et al., 2014).

Thus, a later filter on the ignored signal must actively suppress distracting inputs. We found such a neural filter mechanism (Fig 4C&G) active in a time range which was previously attributed to processing of phonological (Di Liberto et al. 2016, Brodbeck at al. 2018) as well as semantic features (Broderick at al. 2018), which both go beyond basic acoustic properties of speech (Obleser and Eisner 2009). One suggestion of our results is that when phonemes (or even words) of the dominant ignored talker pull bottom-up attention, their representation is actively suppressed at a late stage in order not to impair linguistic representation of the attended talker’s speech.

### Late distractor suppression in a non-auditory, fronto-parietal attention network

Previously, it has been shown that neural selective processing of concurring auditory stimuli is mainly accomplished in auditory cortex, resulting in a ‘clean’ and distraction-invariant representation of the attended talker (Mesgarani and Chang 2012; Zion Golumbic 2013).

Critically, under the adverse SNR of −6 dB, our analysis revealed an enhanced response to the ignored talker in a later time range (i.e., 200-300 ms) consisting of a positive and a negative component (Fig. 3B). The latter is anti-polar to the P2_TRF_ (to the attended talker). This additional component, which we interpret as a signature of active suppression of the ignored talker, involved non-auditory regions, which are part of the fronto-parietal attention or global-demand network (Woolgar et al., 2016), where we found enhanced neural selective processing of the ignored talker.

Under the assumption that such active suppression is costly to the cognitive system, it has been suggested that it is only deployed if necessary (Chait et al., 2010). Neural signatures for active suppression of irrelevant signals during late (~200 ms) AEPs have been examined before (Melara et al., 2002; Chait et al., 2010). Pomper and Chait (2017) related enhanced centro-parietal activity in the theta band (4-7 Hz) to enhanced top-down control. Parietal activity in the theta-band was also found to be inversely related to the delta-band auditory entrainment in superior temporal gyrus (Keitel et al., 2017). Here we show how late top-down, fronto-parietal neural processing of the distracting auditory input is unfolding in time and might facilitate overall selective neural processing.

In earlier studies, researchers highlighted the predominant tracking of the attended talker (Mesgarani and Chang, 2012; Ding & Simon, 2012, Zion Golumbic, 2013, O’Sullivan et al. 2014), emphasizing that a clean representation of the attended talker is key to successful listening. In some contrast to this, previous results shed light on the neural processing of the ignored talker (see also Wöstmann et al., 2017, Olguin et al., 2018). We have shown here that the overall neural selective processing is surprisingly robust against such demanding listening conditions (Ding & Simon, 2013), and that a ‘clean’ or isolated tracking of the ignored talker is at least as essential.

This finding invites some speculation on the neural implementation of attentional filters more generally. On the one hand, a selective neural filter can be solely optimized to let pass relevant features of attended signals. On the other hand, it can be optimized to let pass features of the ignored talker, which might be relevant for suppression at a later stage. In line with earlier studies, we found that the *neural tracking* was dominated by the attended talker (speaking for the first strategy). However, under most demanding listening conditions (i.e., negative SNR), *neural selectivity* was dominated by the ignored talker.

Neural filter mechanisms might thus adapt depending on the listening demand. Follow-up studies should investigate the relationship of such filter adaptation to the concept of listening effort (Rönnberg et al., 2013; McGarrigle et al., 2014): Additional tracking of the ignored talker leads to higher neuro-computational load and might also be related to working memory performance (Rudner et al. 2011).

Within our design, we can only draw limited conclusions on the behavioral impact of the late neural tracking of the ignored talker. This is due to the tradeoff between sufficient behavioral data (e.g., trial-based design) and ecological validity (e.g., presentation of continuous speech; Hamilton and Huth, 2018). Following studies should acquire more fine-grained behavioral data, ideally without losing much of the ecological validity.

Our results show that, within the hierarchy of the central auditory pathways, the cocktail-party problem might look solved or settled at the stage of secondary auditory cortex (Mesgarani & Chang, 2012), but higher-order, attentional networks and their dedicated processing of distracting speech appear key to this solution.

### Conclusions

The present data show how components of the unfolding temporal response function as identified in a forward encoding model of the electroencephalographic signal can reflect distinct neural stages of attentional filtering. These stages contain the initial, attention-independent encoding of acoustic signals; the extraction and amplification of relevant features; and lastly a robust, purely attention-driven selective response to the attended and ignored acoustic signals.

Most consequential to our thinking about attentional filtering in the central auditory system, an active-suppression response to ignored acoustic signals originates from non-auditory, fronto-parietal attentional networks. In sum, with a design closer to real-life listening scenarios, our study provides insight into how selective neural processing of attended speech unfolds and is upheld not only by auditory cortices. Instead, establishing a clean cortical representation of the attended talker as suggested previously hinges on achieving a late suppression of ignored signals, with contributions by regions of the fronto-parietal attention network.

## Methods

### Participants

Eighteen native speakers of German (9 females) were invited from the participant database of the Department of Psychology, University of Lübeck, Germany. We recruited participants who were between 23 and 68 years old at the time of testing (mean: 49, SD: 17), to allow valid conclusions from such a challenging listening scenario to middle-aged and older adults. All reported normal hearing and no histories of neurological disorders. Incomplete data due to recording hardware failure was obtained in four more, initially invited participants. All participants gave informed consent and received payment of 8 €/hour. The study was approved by the local ethics committee of the University of Lübeck.

### Stimuli

The goal of this study was to investigate the selective neural processing of one of two talkers under a continuously varying signal-to-noise ratio (SNR). Here, the signal is a to-be-attended talker and the noise is a to-be-ignored talker. Our study was conducted in a within subject 2 by 3 design (attention by SNR (three levels)).

We selected two audiobooks read by native German speakers, one female (Elke Heidenreich, ‘Nero Corleone kehrt zurück’, read by Elke Heidenreich) and one male (Yuval Noah Harari, ‘Eine kurze Geschichte der Menschheit’, read by Jürgen Holdorf). The following steps of stimulus preparation were done using custom code written in MATLAB (Version 2017a; *Mathworks Inc., Natick*, M/A). Sequences of silence longer than 500 ms were truncated to 500 ms to avoid long parts of silence (O’Sullivan et al., 2014). The first hour of each audiobook was selected for further preparation. The first 30 minutes of each audiobook served as the to-be-attended and the rest served as the to-be-ignored speech, such that all subjects could attend both stories from the beginning and attended (and ignored) both the female and the male voice the same amount of time.

The identical mixture of the attended and ignored talker was presented on both ears, resulting in a concurrent listening scenario without any spatial cue (i.e. diotic, Fig. 1 A). Hence, the only cues available for talker segregation consisted in the spectro-temporal features of the talkers, such as pitch, formants, and amplitude modulation.

The SNR was modulated symmetrically around 0 dB. An SNR of 0 dB refers to concurrent talker signals with a matched long-term root-mean-square (rms) amplitude as used previously in numerous studies (e.g. Power et al., 2012; O’Sullivan et al., 2014; Mirkovic et al., 2015). Coming from an SNR of 0dB, the SNR was either increased to +6 dB by raising the sound pressure level (SPL) of the to-be-attended talker by 6 dB or decreased to −6 dB by raising the SPL of the to-be-ignored talker by 6 dB. Thus, the talkers were either *balanced* (Fig. 1B, black) or one of the talkers was *dominant* (Fig. 1B, purple) and the other was *non-dominant* (Fig. 1B, grey). The particular SNR-range (−6 to +6 dB) was chosen to create a challenging but at the same time solvable listening task. Even if an SNR of-6 dB is rare in real-life listening scenarios (Smeds et al., 2015), the neural tracking of attended speech has been reported as intact at SNRs as low as −6 dB (Ding and Simon, 2013). However, speech perception (number of words repeated correctly) of normal hearing subjects starts to suffer around an SNR < 0 dB and the speech-reception threshold (i.e. 50% correct) usually lies between −5 and 0 dB (Pichora-Fuller et al., 1995, Bentler et al., 2004).

As building blocks for SNR modulation, we created a sample of plateaus (i.e., constant SNR of −6, 0 or +6 dB) and ramps (i.e., transition between plateaus). The length of plateaus was uniformly distributed between 5 and 9 seconds in discrete steps of one second. The ramps were linear interpolations between SNRs with the length uniformly distributed between 1 and 5 seconds in discrete steps of one second. The length distributions of plateaus and ramps were kept uniform within each talker and within their assignments as being attended or ignored. We concatenated plateaus via ramps such that a 0 dB plateau was either followed by a +6 dB or a −6 dB, whereas a +6 dB or a −6 dB plateau were always followed by a 0 dB plateau via a respective ramp. Randomly varying SNR time courses were created for each subject individually in order to avoid systematic overlap between the SNR modulation and the audiobooks. Stimulus material was cut into twelve blocks, which resulted in an average block length of five minutes. Sound files were created with a sampling rate of 44.1 kHz and a 16-bit resolution. The experiment was implemented in the software *Presentation (Neurobehavioural Systems)*.Stimuli were presented via headphones (Sennheiser HD25).

### Task

The twelve blocks were presented such that subjects were instructed to attend to the female or to the male talker in an alternating fashion. After instruction before each block (i.e. attend to female or attend to male), subjects were asked to start the stimulus presentation by a button press, which enabled the participants to take a break between blocks. During listening, subjects were asked to fixate a cross presented on the screen in order to reduce eye movement.

Every other block, the stories picked up at the point it ended two blocks before. After each block, subjects were asked to rate the difficulty of maintaining attention by mouse-clicking on a continuous color bar ranging from red (difficult) to green (easy). For later analysis, the continuous color bar was discretized into ten segments (1 = difficult, 10 = easy). Subsequently, participants were asked to answer four multiple-choice questions concerning the content of the to-be-attended audiobook. The average rating of difficulty was neither significantly correlated with the number of questions correctly answered (Pearson’s r = 0.1, p = 0.73), nor with participants’age (Pearson’s r = −0.17, p = 0.51). Furthermore, we found no significant correlation of the number of correctly answered questions with age (Pearson’s r = −0.11, p = 0.65).

### Data acquisition and preprocessing

EEG was recorded with 64 electrodes *Acticap* (*Easycap, Herrsching, Germany*) connected to an *ActiChamp*(*Brain Products, Giiching, Germany*) amplifier. EEG signals were recorded with the software *Brain Vision Recorder* (*Brain Products*) at a sampling rate of 1 kHz. Impedances were kept below 10 kΩ. Electrode TP9 (left mastoid) served as reference during recording.

The EEG data were pre-processed in *MATLAB (2017a)* using both the *Fieldtrip*-toolbox (version: 20170321; Oostenveld et al., 2011) and custom written code. The EEG data were re-referenced to the average of the electrodes TP9 and TP10 (left and right mastoids) and resampled to *f_s_* = 125 Hz. The continuous EEG data were highpass-filtered at *f_c_* = 1 Hz and lowpass-filtered at *f_c_* = 30 Hz (two-pass Hamming window FIR, filter order: 3f_s_/f_c_).

From the continuous EEG data, we extracted the parts during which the twelve blocks of audiobooks were presented (see above). For every subject, we applied independent component analysis (ICA; Makeig et al., 2004) on the concatenated data of the twelve blocks and manually rejected components that were clearly related to eye movements, eye blinks, muscle artifacts, heartbeat as well as singe-channel noise. On average, 26 of 62 components (SD: 7.3) were rejected.

For further analysis, we lowpass-filtered the data again at f_c_ = 10 Hz (two-pass Hamming window FIR, filter order: 3f_s_/f_c_), which assured that the amplitudes at all frequencies up to 8 Hz were not reduced. Previously, neural activity phase-locked to the envelope was only found up to a frequency of approximately 8 Hz (Zion Golumbic et al., 2013; Ding et al., 2014). We could confirm this finding by incrementally raising the cutoff frequency, which didn’t change the morphology of the TRFs (see below) but only decreased the prediction accuracy due to the interference of nonphase-locked neural activity and external noise in higher frequencies.

### Extraction of envelope onsets

A temporal representation of the acoustic onsets, further called envelope onsets, was extracted from the presented speech signals (Fiedler et al., 2017). Those representations later served as regressors to model neural responses to the talkers (see below). First, we extracted an auditory spectrogram containing 128 spectrally resolved sub-band envelopes of the speech signals logarithmically spaced between approximately 90 and 4000 Hz using the *NSL* toolbox (Chi et al., 2005). Second, the auditory spectrogram was summed up across frequencies, which resulted in broadband temporal envelopes of the audiobooks. Taking the derivative of the envelope and zeroing all values smaller than zero (Hertrich et al., 2012) returned the envelope onsets, which only contain positive values at time periods of an increasing envelope, as can be found at acoustic onsets (Fig. 1C).

Using the envelope onsets as regressor does not imply that we only modeled the encoding of acoustic onsets. Every onset is followed by a peak in the speech envelope (Fig. 1C), which is then again followed by an offset and the next onset and so forth, resulting in a high autocorrelation between those features. Nevertheless, onsets are the earliest feature that could possibly evoke a neural response (Picton, 2013). The latency of modeled responses to envelope onsets (compared to envelopes) was found to be most similar to conventional ERPs (Fiedler et al., 2017, supplemental material).

### Estimation of temporal response functions

We applied an established method to estimate a linear forward (encoding) model (Lalor et al., 2009; Crosse et al., 2016). The model contains temporal response functions (TRFs), which are estimations of the neural response to a continuously varying stimulus feature. In our case, this stimulus feature is the envelope onsets (see above) of both, the attended and the ignored talker. Based on the assumption that every sample in the EEG signal *r(t)* is the superposition of neural responses to past onsets and thus can be expressed for one talker by a convolution operation:

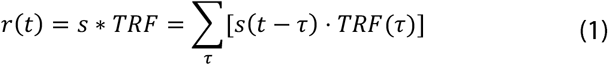

where *s(t)* is the envelope onsets, TRF is the temporal response function that describes the relationship between s and r over a range of time lags r (Fig. 1C). The TRF contains a weight for each time lag *τ*. We investigated time lags in the range from −100 to 500 ms. In order to obtain the β-weights of the TRF to both talkers contained in the matrix *G_TRF_*, ridge regression (Hoerl and Kennard, 1970) was applied, which can be expressed in the linear algebraic form:

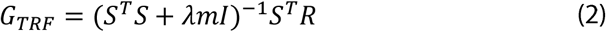

where *S* is matrix containing the onset envelopes of both the attended and ignored talker and its sample-wise time lagged replications, *R* contains the measured EEG signal, λ is the ridge parameter for regularization, the scalar *m* is the mean of the trace of S^T^S (Biesmans et al., 2016) and *I* is the identity matrix. The optimal ridge parameter λ was estimated according to Fiedler et al. (2017) and was set to λ= 10.

TRFs were estimated on a trial-by-trial basis, where trial refers to a part (e.g. a plateau of +6 dB) of certain length cut from the continuous stimulus and the respective EEG data. For the subsequent analysis, we subdivided the data in two ways: First, to get a general estimate of the model’s ability to dissociate between attended and ignored talkers, we cut the data into one-minute trials, resulting in trial lengths comparable to previous studies (O’Sullivan et al., 2014; Mirkovic et al., 2015; Biesmans et al., 2016; Fiedler et al., 2017). This resulted in 60 trials per subject. Second, we cut the data based on the applied SNR modulation, which resulted in three groups of trials: −6 dB, 0 dB and +6 dB. To use the entire recording, the data were cut at the time points where ramps of the SNR time courses either crossed −3 dBor+3 dB (Fig. 1B). This resulted in 180 trials ofOdB and 90 trials of-6 and +6 dB, respectively. The average length of those trials was 10 seconds (i.e. average length of a plateau (7 seconds) and average length of two halves of a ramp (2×1.5 seconds)). In order to balance the number of trials across SNRs, 90 trials from 0 dB were randomly drawn from the 180 trials of every subject. During the analysis, we contrasted TRFs not only within conditions, but also contrasted the TRFs to the talkers within their role of being dominant (Fig 2B, purple; attended under SNR =+6 dB, ignored under SNR = −6 dB) or non-dominant (Fig 2B, grey; attended under SNR = −6 dB, ignored under SNR =+6 dB). We will use those terms and schematic bar graphs (Fig. 1B) throughout the entire article.

### Statistical analysis on temporal response functions

To extract significant spatio-temporal deflections in the TRFs at an SNR of 0 dB, we applied a two-level statistical analysis (two-level cluster-test; e.g. Obleser et al., 2012). At the single-subject level, we used one-sample t-tests to test the TRF to the attended, the ignored as well as the attended-ignored difference against zero. Resulting t-values were transformed to z-scores. At the group level, the deflection of z-scores from zero was tested by a cluster-based permutation one-sample t-test (Maris and Oostenveld, 2007), which clusters t-values with p-values < 0.001 of adjacent time-electrode bins (with a minimum of 4 neighboring electrodes). The extracted cluster is compared to 4,000 clusters drawn randomly from the data by permuting condition labels. The resulting cluster p-value reflects the relative number of Monte Carlo iterations in which the summed t-statistic of the observed cluster is exceeded. This contrast indicates how components of the TRF are generally affected by attention under balanced conditions.

In a second step, the identical cluster-based permutation test was applied to obtain significant differences between the TRFs depending on whether a talker was dominant or non-dominant. This contrast was separately computed for the attended an ignored talker and it indicates, how the TRFs are affected by changing SNR.

In a third step, the difference between the TRFs to the attended and ignored talker were contrasted separately for *dominant* and *non-dominant* talkers. This contrast describes how attention affects the TRF to a dominant talker (easy-to-attend, hard to ignore) or a non-dominant talker (hard-to-attend, easy-to-ignore), respectively.

For illustration of the neural responses, we averaged single-subject TRF β-weights across channels of interest. Channels of interest were defined as the channels being part of both significant clusters found in the attended-ignored difference between TRFs under a balanced SNR of 0 dB (Fig. 2B). The 95%-confidence-bands were obtained by bootstrapping (Efron, 1979) across the averaged TRFs of all subjects, using 4,000 iterations.

### Neural tracking and neural selectivity

To disentangle bottom-up and top-down effects, we investigated the TRFs based on two measures: *neural tracking* and *neural selectivity*. While *neural tracking* is a measure of how strongly a talker is encoded in the EEG (irrespective of attention), *neural selectivity* is a measure of how differential (i.e., attended vs. ignored) those representations are due to the impact of selective attention.

As a base for those two measures, we followed the forward method of predicting EEG signals and comparing those to the measured EEG signal, as described in detail by in Fiedler et al. (2017). In a leave-one-out fashion, we predicted EEG signals of a single trial contained in Ř following the equation:

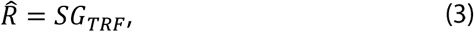

where S is the matrix containing the onset envelopes and G_TRF_ is the matrix containing the trained TRFs.

*Neural tracking* was defined as the Pearson-correlation coefficient between the predicted and recorded EEG signals using the estimated TRFs (see above).

*Neural selectivity* was defined as the percentage of trials the TRFs could successfully identify a talker as being attended or ignored. Therefore, two different EEG signals were predicted per trial (Eq. 3), the first representing a talker being attended and the second representing the same talker being ignored. While one of the EEG signals is representing the task instruction (i.e., attend the to-be-attended talker; ignore the to-be-ignored talker), the other EEG signal represents the alternative (i.e. attending the to-be-ignored talker; ignoring the to-be-attended talker). We calculated the Pearson correlations for both predicted EEG signals with the measured EEG signal (Fiedler et al., 2017). Talker identification was successful if the EEG signal referring to the task instruction yielded higher correlation. Note that during unbalanced SNRs (i.e., −6 dB & +6 dB), the alternative EEG signal was predicted based on the TRFs estimated on the opposite SNR (e.g., under an SNR of +6 dB, the alternative to attending the to-be-attended talker (*dominant*) is ignoring the to-be-ignored talker under an SNR of −6 dB).

Since this is a forward model approach, *neural tracking* and *neural selectivity* were obtained at every single EEG channel (Crosse et al., 2016). Likewise, both measures were obtained at the source level at every single voxel. We split up the prediction by either using only the prediction of the to-be-attended, only the prediction of the to-be-ignored or the sum of both predictions, such that the talker-specific contribution to *neural tracking* (*neural selectivity*) could be compared to the overall *neural tracking* (*neural selectivity*).

In order to evaluate the unfolding of *neural tracking* and *neural selectivity*over TRF time lags, we used a sliding-window of time lags (size: 48 ms, 6 samples) with an overlap of 24 ms (3 samples) for the prediction. For every position of the window, *neural tracking* and *neural selectivity* were calculated (see above).

In advance of any arithmetic operation on *neural tracking*, the underlying Pearson-correlation coefficients were fisher-z transformed. Accordingly, *neural selectivity* (i.e., percentage correct) was logit-transformed.

### Source localization

To further trace the origin of effects observed in sensor space, we applied LCMV-beamforming (Drongelen et al., 1994; Van Veen et al., 1997) to obtain source-activity time courses in single voxels of the brain. Using a standard template brain from Fieldtrip/SPM (Montreal Neurological Institute) together with the *Acticap* electrode layout, leadfields were calculated with a grid resolution of 10 mm. Individual LCMV-filter weights were obtained using 5% regularization. The continuous time-domain EEG data were projected to source space, resulting in three source activity time courses (X-Y-Z) per voxel. In order to obtain a single time course for each voxel, the direction of highest variance was determined by principal component analysis and used for further analysis. All further processing steps in source space were done analogously to sensor space EEG data.

### Appendix

**Figure S1:**
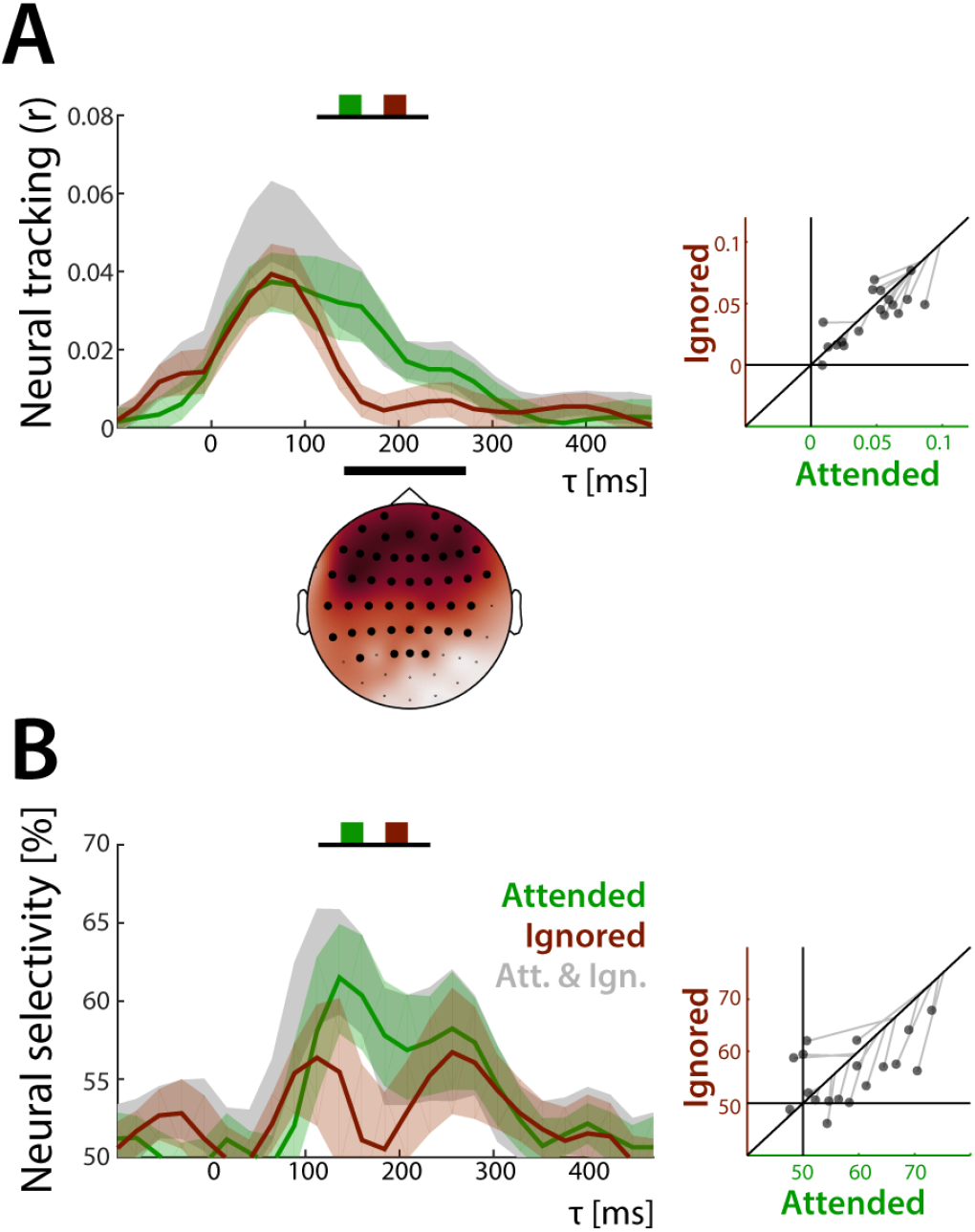
Unfolding of neural tracking and neural selectivity under the balanced SNR of 0 dB. Neural tracking and neural selectivity were estimated based on the extracted TRFs to the attended (green), the ignored (red) as well as both talkers (grey). Confidence bands (95%) were obtained by bootstrapping. Highlighted channels (topographic maps) are part of a significant cluster. **A)** Neural tracking across all time lags (−100-500 ms). Scatterplots (bottom) show single-subject data averaged across channels of interest. Grey lines indicate overall neural tracking of both talkers at the 45°-line. **B)** Neural selectivity across all time lags (−100-500 ms). Scatterplots (bottom) show single-subject data averaged across channels of interest. Grey lines indicate overall neural selectivity of both talkers at the 45°-line.

#### Extraction of peak latencies and peak amplitude

In order to disentangle amplitude- and latency-effects, we extracted peak latency and peak amplitude for every subject and every component.

Peak latencies were defined as the time lag of the maximum or minimum within a certain time interval (P1_TRF_: 0−100 ms; N1_TRF_: 100-200 ms; P2_TRF_: 200-350 ms) of the subject- and SNR-specific TRF. The peak amplitudes were defined as the β-weights at the respective peak latencies. Please note that a reliable extraction of a P2 component was only possible in the TRF to the attended talker, whereas a reliable estimation of the N2 component was only achieved in the TRF to the dominant ignored talker under −6.

The main effects and interactions of attention and SNR on both the peak latency and peak amplitude were investigated by a repeated-measures ANOVA. Reported p-values were obtained with Greenhouse-Geisser-corrected degrees of freedom.

#### SNR-induced TRF latency shift

The contrasted TRFs between the *dominant* and *non-dominant* attended talker (Fig. 3A) showed significant differences during three time-lag intervals, first at around 20 ms (8-24 ms, p = 0.004), a second around 100 ms (80-128 ms, p = 2×10^4^) and a third around 200 ms (176-224 ms, p = 0.001). These differences occurred in the transition between components (P1_TRF_ to N1_TRF_, and N1_TRF_ to P2_TRF_). This was consistent with the visual impression of the TRFs being similar in morphology yet delayed whenever a talker was less dominant. The TRFs to the ignored talker (Fig. 3B) also suggest such an SNR-related delay, even if no comparable significant differences were observed at the transitions between components. Nevertheless, the individual peak latencies showed a main effect of SNR for the P1 TRF (the more dominant the earlier; Fig. S2, F_1.64,27.87_ = 21.67, p = 0.006×10^−4^), but also a main effect of attention (earlier if attended, F_1,17_ = 13.73, p = 0.002). No interaction between SNR and attention was found (F_1.61,27.42_ = 1.93, p = 0.171; see appendix for more details).

**Figure S2:**
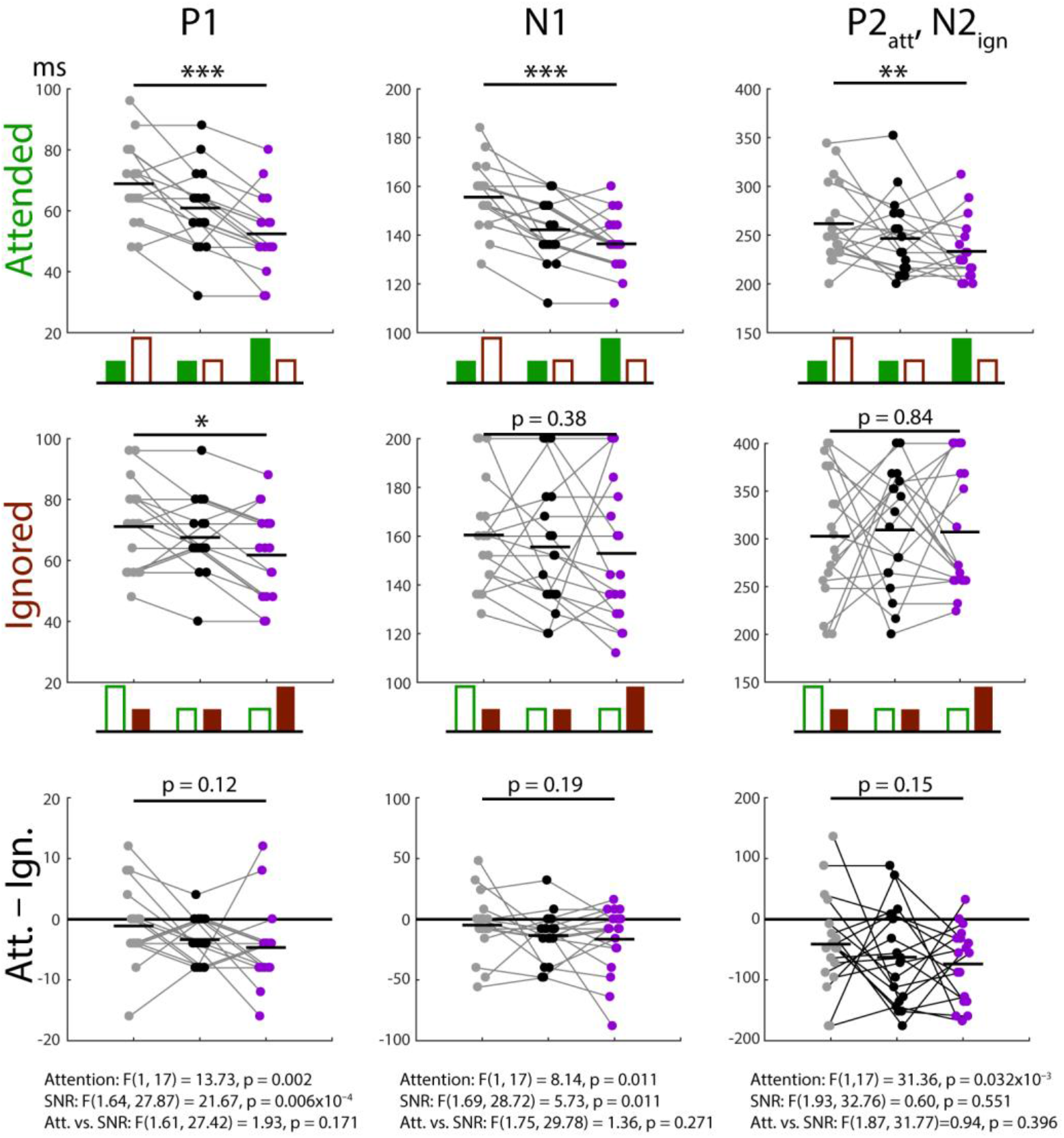
Peak latencies extracted from TRFs of single subjects for dominant (purple), balanced (black) and non-dominant talkers (attended and ignored).

**Figure S3:**
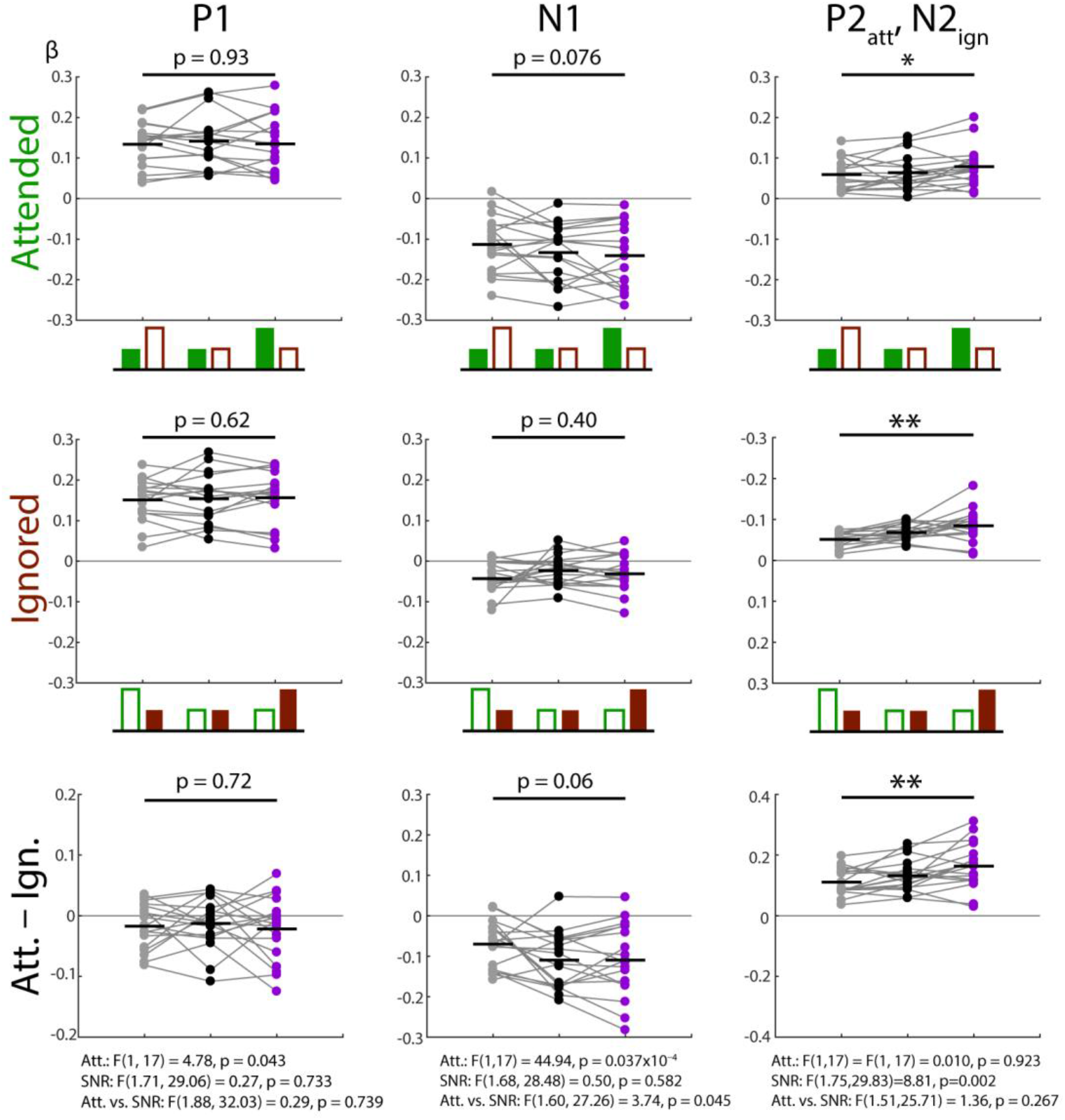
Peak amplitudes extracted from TRFs of single subjects for dominant (purple), balanced (black) and non-dominant talkers (attended and ignored).

## Acknowledgments

Research was supported by the European Research Council (ERC-CoG-2014 646696 to JO) and the Oticon Foundation (NEUROCHAT).

## References

Bentler RA, Palmer C, Dittberner AB. 2004. Hearing-in-Noise: Comparison of Listeners with Normal and (Aided) Impaired Hearing. J Am Acad Audiol. 15:216–225.

Biesmans W, Das N, Francart T, Bertrand A. 2016. Auditory-inspired speech envelope extraction methods for improved EEG-based auditory attention detection in a cocktail party scenario. {IEEE} Trans Neural Syst Rehabil Eng. 4320.

Brodbeck C, Hong LE, Simon JZ. 2018. Transformation from auditory to linguistic representations across auditory cortex is rapid and attention dependent for continuous speech. bioRxiv.

Broderick MP, Anderson AJ, Liberto GM Di, Crosse MJ, Edmund C. 2018. Electrophysiological correlates of semantic dissimilarity reflect the comprehension of natural, narrative speech. Curr Biol. 28:803–809.

Chait M, Cheveigné A De, Poeppel D, Simon JZ. 2010. Neuropsychologia Neural dynamics of attending and ignoring in human auditory cortex. Neuropsychologia. 48:3262–3271.

Cherry EC. 1953. Some Experiments on the Recognition of Speech, with One and with Two Ears. J Acoust Soc Am. 25:975–979.

Chi T, Ru P, Shamma SA. 2005. Multiresolution Spectrotemporal Analysis of Complex Sounds. J Acoust Soc Am. 118:887–906.

Combrisson E, Jerbi K. 2015. Exceeding chance level by chance: The caveat of theoretical chance levels in brain signal classification and statistical assessment of decoding accuracy. J Neurosci Methods. 1–11.

Crosse MJ, Di Liberto GM, Bednar A, Lalor EC. 2016. The Multivariate Temporal Response Function (mTRF) Toolbox: A MATLAB Toolbox for Relating Neural Signals to Continuous Stimuli. Front Hum Neurosci. 10:604.

Davis MH, Johnsrude IS. 2003. Hierarchical Processing in Spoken Language Comprehension. 23:3423–3431.

Di Liberto GM, O’Sullivan JA, Lalor EC. 2015. Low-frequency cortical entrainment to speech reflects phoneme-level processing. Curr Biol. 25:2457–2465.

Ding N, Chatterjee M, Simon JZ. 2014. Robust cortical entrainment to the speech envelope relies on the spectro-temporal fine structure. Neuroimage. 88:41–46.

Ding N, Simon JZ. 2012. Neural coding of continuous speech in auditory cortex during monaural and dichotic listening. J Neurophysiol. 107:78–89.

Ding N, Simon JZ. 2013. Adaptive temporal encoding leads to a background-insensitive cortical representation of speech. J Neurosci. 33:5728–5735.

Drongelen W Van, Yuchtman M, Veen BD Van, Huffelen AC Van. 1994. A Spatial Filtering Technique to Detect and Localize Multiple Sources in the Brain. Brain Topogr. 9:39–49.

Efron B. 1979. Bootstrap methods: Another look at the jackknife. Ann Stat. 7:1–26.

Fiedler L, Wöstmann M, Graversen C, Brandmeyer A, Lunner T, Obleser J. 2017. Single-channel in-ear-EEG detects the focus of auditory attention to concurrent tone streams and mixed speech. J Neural Eng. 14.

Fuglsang SA, Dau T, Hjortkjær J. 2017. NeuroImage Noise-robust cortical tracking of attended speech in real-world acoustic scenes. Neuroimage. 156:435–444.

Giordano BL, Ince RAA, Gross J, Schyns PG, Panzeri S, Kayser C. 2017. Contributions of local speech encoding and functional connectivity to audio-visual speech perception. Elife. 6:1–27.

Hamilton LS, Huth AG. 2018. The revolution will not be controlled: natural stimuli in speech neuroscience. Lang Cogn Neurosci. 1–10.

Hertrich I, Dietrich S, Trouvain J, Moos A, Ackermann H. 2012. Magnetic brain activity phase-locked to the envelope, the syllable onsets, and the fundamental frequency of a perceived speech signal. Psychophysiology. 49:322–334.

Hoerl AE, Kennard RW. 1970. Ridge Regression: Biased Estimation for Nonorthogonal Problems. Technometrics. 12:55–67.

Horton C, Zmura MD, Srinivasan R. 2013. Suppression of competing speech through entrainment of cortical oscillations. J Neurophysiol. 109:3082–3093.

Kaya EM, Elhilali M. 2017. Modelling auditory attention. Phil Trans R Soc B. 372.

Kayser SJ, Ince RAA, Gross J, Kayser C. 2015. Irregular Speech Rate Dissociates Auditory Cortical Entrainment, Evoked Responses, and Frontal Alpha. J Neurosci. 35:14691–14701.

Keitel A, Ince RAA, Gross J, Kayser C. 2017. NeuroImage Auditory cortical delta-entrainment interacts with oscillatory power in multiple fronto-parietal networks. Neuroimage. 147:32–42.

Kong Y-Y, Somarowthu A, Ding N. 2015. Effects of Spectral Degradation on Attentional Modulation of Cortical Auditory Responses to Continuous Speech. J Assoc Res Otolaryngol. 16:783–796.

Kong YY, Mullangi A, Ding N. 2014. Differential modulation of auditory responses to attended and unattended speech in different listening conditions. Hear Res. 316:73–81.

Lalor EC, Power AJ, Reilly RB, Foxe JJ. 2009. Resolving Precise Temporal Processing Properties of the Auditory System Using Continuous Stimuli. J Neurophysiol. 102:349–359.

Makeig S, Debener S, Onton J, Delorme A. 2004. Mining event-related brain dynamics. Trends Cogn Sci. 8:204–210.

Maris E, Oostenveld R. 2007. Nonparametric statistical testing of EEG- and MEG- data. J Neurosci Methods. 164:177–190.

Mcgarrigle R, Munro KJ, Dawes P, Stewart AJ, David R, Barry JG, Amitay S. 2014. Listening effort and fatigue: What exactly are we measuring? Int J Audiol. 53:433–445.

Melara RD, Rao A, Tong Y. 2002. The duality of selection: Excitatory and inhibitory processes in auditory selective attention. J Exp Psychol. 28:279–306.

Mesgarani N, Chang EF. 2012. Selective cortical representation of attended speaker in multi-talker speech perception. Nature. 485:233–236.

Mirkovic B, Debener S, Jaeger M, Vos M De. 2015. Decoding the attended speech stream with multi-channel EEG: implications for online, daily-life applications. J Neural Eng. 12:46007.

O’Sullivan JA, Power AJ, Mesgarani N, Rajaram S, Foxe JJ, Shinn-Cunningham BG, Slaney M, Shamma SA, Lalor EC. 2014. Attentional Selection in a Cocktail Party Environment Can Be Decoded from Single-Trial EEG. Cereb Cortex. 25:1697–1706.

Obleser J, Eisner F. 2009. Pre-lexical abstraction of speech in the auditory cortex. Trends Cogn Sci. 13:14–19.

Obleser J, Wöstmann M, Hellbernd N, Wilsch A, Maess B. 2012. Adverse Listening Conditions and Memory Load Drive a Common Alpha Oscillatory Network. J Neurosci. 32:12376–12383.

Olguin A, Bekinschtein TA, Bozic M. 2018. Neural Encoding of Attended Continuous Speech under Different Types of Interference. J Cogn Neurosci. (in Print).

Oostenveld R, Fries P, Maris E, Schoffelen J. 2011. FieldTrip: Open Source Software for Advanced Analysis of MEG, EEG, and Invasive Electrophysiological Data. Comput Intell Neurosci. 2011.

Petersen EB, Wöstmann M, Obleser J, Lunner T. 2016. Neural tracking of attended versus ignored speech is differentially affected by hearing loss. J Neurophysiol. 117:18–27.

Pichora-Fuller MK, Schneider BA, Daneman M. 1995. How young and old listen to and remember speech in noise. J Acoust Soc Am. 91:593–608.

Picton T. 2013. Hearing in Time: Evoked Potential Studies of Temporal Processing. Ear Hear. 34:385–401.

Pomper U, Chait M. 2017. The impact of visual gaze direction on auditory object tracking. Sci Rep. 7:1–16.

Power AJ, Foxe JJ, Forde EJ, Reilly RB, Lalor EC. 2012. At what time is the cocktail party? A late locus of selective attention to natural speech. Eur J Neurosci. 35:1497–1503.

Rönnberg J. 2013. The Ease of Language Understanding (ELU) model: theoretical, empirical, and clinical advances. Front Syst Neurosci. 7:1–17.

Rudner M, Rönnberg J, Lunner T. 2011. Working Memory Supports listening in Noise for Persons with Hearing Impairment. J Am Acad Audiol. 22:156–167.

Smeds K, Wolters F, Rung M. 2015. Estimation of Signal-to-Noise Ratios in Realistic Sound Scenarios. J Am Acad Audiol. 196:183–196.

Veen BD Van, Drongelen W Van, Yuchtman M, Suzuki A. 1997. Localization of Brain Electrical Activity via Linearly Constrained Minimum Variance Spatial Filtering. IEEE Trans Biomed Eng. 44:867–880.

Wang Y, Zhang J, Ding N, Zou J, Luo H. 2018. Prior Knowledge Guides Speech Segregation in Human Auditory Cortex. Cereb Cortex. bhy052:1–11.

Waschke L, Wöstmann M, Obleser J. 2017. States and traits of neural irregularity in the age-varying human brain. 1–12.

Willmore BDB, Cooke JE, King AJ. 2014. Hearing in noisy environments: noise invariance and contrast gain control. J Physiol. 16:3371–3381.

Woolgar A, Jackson J, Duncan J. 2016. Coding of Visual, Auditory, Rule, and Response Information in the Brain: 10 Years of Multivoxel Pattern Analysis. J Cogn Neurosci. 28:1433–1454.

Wöstmann M, Fiedler L, Obleser J. 2016. Tracking the signal, cracking the code: Speech and speech comprehension in non-invasive human electrophysiology. Languange, Cogn Neurosci. 1–15.

Wöstmann M, Lim S, Obleser J. 2017. The Human Neural Alpha Response to Speech is a Proxy of Attentional Control. Cereb Cortex. 27:3307–3317.

Zion Golumbic EM, Ding N, Bickel S, Lakatos P, Schevon CA, McKhann GM, Goodman RR, Emerson R, Mehta AD, Simon JZ, Poeppel D, Schroeder CE. 2013. Mechanisms underlying selective neuronal tracking of attended speech at a “cocktail party.” Neuron. 77:980–991.

